# *In silico* predictions of the hepatic metabolic clearance in humans for 10 drugs with highly variable *in vitro* pharmacokinetics

**DOI:** 10.1101/2023.04.01.535222

**Authors:** Urban Fagerholm

## Abstract

Challenges/problems for *in vitro* methodologies for prediction of human clinical pharmacokinetics include inter- and intra-laboratory variability, and common occurance of high limits of quantification, low recovery, low parameter validity and low reproducibility. In this study, 10 drugs with substantial differences in human hepatocyte intrinsic metabolic clearance (CL_int_) and fraction unbound in plasma (f_u_) between laboratories were selected. The average and maximum ratios between highest and lowest reported predicted *in vivo* hepatic metabolic clearance (CL_H_) for the drugs were 529- and 2436-fold, respectively. The *in vivo* CL_H_ was predicted using *in vitro* CL_int_ and f_u_ data from the various highly sources and using our *in silico* methodology. The main aim was to compare the predictive accuracies for the *in vitro* and *in silico* methodologies. Prediction errors for *in vitro* methodology ranged from 1.1-to 578-fold, with an average of 150-fold for lowest predicted estimates and 16-fold for highest predicted estimates. The *in vitro* based predictions produced 36-to 38-fold higher average and maximum prediction errors than the *in silico* methodology, respectively. Mean and maximum *in silico* prediction errors were 4.2- and 15-fold, respectively, which is consistent with earlier results. In contrast to the *in vitro* methodology the *in silico* models did not predict high hepatic extraction ratio for drugs with low CL_H_. Overall, the *in silico* method clearly outperformed *in vitro* data for prediction of CL_H_ in man for 10 drugs with large interlaboratory variability.

## INTRODUCTION

Challenges/problems for *in vitro* methodologies for prediction of human clinical pharmacokinetics (PK) include inter- and intra-laboratory variability, and common occurance of high limits of quantification (such as for conventional hepatocyte and microsome assays), low recovery (such as for the Caco-2 assay), low parameter validity (such as for aqueous solubility (S) *vs in vivo* dissolution potential) and low reproducibility.^1–8^

Ten physico-chemically and pharmacokinetically different drugs with large interlaboratory variability of intrinsic metabolic hepatic clearance (CL_int_) and unbound fraction (f_u_) are shown in Table 1 - atenolol, cimetidine, desipramine, diazepam, diclofenac, diltiazem, gemfibrozil, midazolam, naloxone and naproxen.^1–10^ The average and maximum ratios between highest and lowest reported predicted *in vivo* CL_int_ for these are 43- and 138-fold, respectively. Corresponding ratios for measured fu are 13- and 50-fold, respectively. For gemfibrozil, diclofenac and naproxen, CL_int_ • f_u_ differs 3442-, 2842- and 1654-fold depending of choice of sources for CL_int_ and f_u_, respectively.

**Table 1.** Predicted *in vivo* CL_int_, measured fu and ratios between upper and lower estimates for the 10 selected drugs.

| In vitro data and predictions |  |  |  |  |  |  |
| --- | --- | --- | --- | --- | --- | --- |
| Drug | Low CL<sub>int</sub><br>mL/min | High CL<sub>int</sub><br>mL/min | Ratio<br>-fold | Low f<sub>u</sub> | High f<sub>u</sub> | Ratio |
| Atenolol | <200 | 1719 | >8,6 | 0,49 | 0,97 | 2,0 |
| Cimetidine | 227 | 13850 | 61 | 0,52 | 0,87 | 1,7 |
| Desipramine | 770 | 12390 | 16 | 0,093 | 0,16 | 1,7 |
| Diazepam | 51 | 1701 | 33 | 0,0080 | 0,0360 | 4,5 |
| Diclofenac | 505 | 28700 | 57 | 0,0020 | 0,100 | 50 |
| Diltiazem | 2345 | 11690 | 5,0 | 0,0244 | 0,25 | 10 |
| Gemfibrozil | 90 | 12390 | 138 | 0,0020 | 0,050 | 25 |
| Midazolam | 777 | 10710 | 14 | 0,0170 | 0,0720 | 4,2 |
| Naloxone | 1114 | 41720 | 37 | 0,175 | 0,722 | 4,1 |
| Naproxen | 16 | 987 | 62 | 0,0007 | 0,0180 | 27 |
|  |  | Minimum | 5,0 |  |  | 1,7 |
|  |  | Median | 35 |  |  | 4,4 |
|  |  | Mean | 43 |  |  | 13 |
|  |  | Maximum | 138 |  |  | 50 |

For both CL_int_ and f_u_ the ratio between highest and lowest *in vitro* values tend to increase with number of measurements/sources (R^2^-values of 0.36 and 0.18 for n *vs* log CL_int_ and f_u_, respectively).^1,3^ Based on these correlations, the ratio between highest and lowest CL_int_ for a compound increases by 10-fold when increasing the number of measurements from 1 to 10. A 3.1-fold increase is expected for f_u_. Thus, for every additional source there is a higher chance to find a deviating (but not necessarily less accurate) predicted estimate. Such variability/uncertainty is not an issue with *in silico* prediction methods.

We have developed and validated *in silico* methodology for prediction of prediction of human clinical PK, including the ANDROMEDA by Prosilico software.^3,4,7,8,11–13^ In a major benchmarking study ANDROMEDA outperformed laboratory methods in predictive accuracy and range.^8^ Our *in silico* methodology has also outperformed animal models for prediction of volume of distribution and oral bioavailability and hepatocytes for prediction of CL_int_.^4,11,13^ PK-predictions from ANDROMEDA have also been approved by the German authority BfArM for use and main preclinical PK-source in clinical trial applications.

The main objective of this study was to compare the accuracy of our previously developed *in silico* methodology and the laboratory data-based methodology to predict the human CL_H_ of atenolol, cimetidine, desipramine, diazepam, diclofenac, diltiazem, gemfibrozil, midazolam, naloxone and naproxen.

## METHODOLOGY

### Compound and data selection

Atenolol, cimetidine, desipramine, diazepam, diclofenac, diltiazem, gemfibrozil, midazolam, naloxone and naproxen were selected as compounds as these have demonstrated large variability of predicted *in vivo* CL_int_ and measured *in vitro* f_u_ between laboratories/sources (Tables 1 and 2).^1–3,9,10^

**Table 2.** Predicted and observed/measured *in vivo* CL_H_, ratios between upper and lower estimates and estimated minimum and maximum prediction errors for the 10 selected drugs.

| In vitro data and predictions |  |  |  |  |  | In vivo data |
| --- | --- | --- | --- | --- | --- | --- |
| Drug | Low CL<sub>H</sub><br>mL/min | High CL<sub>H</sub><br>mL/min | Ratio<br>-fold | CL<sub>H</sub> pred<br>low error | CL<sub>H</sub> pred<br>high error | In vivo CL<sub>H</sub><br>mL/min |
| Atenolol | <92 | 790 | >8,6 | <4,6 | 39 | 20 |
| Cimetidine | 109 | 1334 | 12 | 7,8 | 95 | 14 |
| Desipramine | 68 | 854 | 13 | 11 | 1,1 | 756 |
| Diazepam | 0,41 | 59 | 144 | 65 | 2,2 | 27 |
| Diclofenac | 1,01 | 985 | 976 | 240 | 4,1 | 243 |
| Diltiazem | 55 | 991 | 18 | 4,8 | 3,8 | 264 |
| Gemfibrozil | 0,18 | 438 | 2436 | 578 | 4,2 | 104* |
| Midazolam | 13 | 509 | 39 | 28 | 1,4 | 371 |
| Naloxone | 173 | 1429 | 8,3 | 3,1 | 2,7 | 535* |
| Naproxen | 0,011 | 18 | 1634 | 558 | 2,9 | 6* |
|  |  | Minimum | 8,3 | 3,1 | 1,1 |  |
|  |  | Median | 28 | 20 | 3,3 |  |
|  |  | Mean | 529 | 150 | 16 |  |
|  |  | Maximum | 2436 | 578 | 95 |  |

\*Extrahepatic and renal elimination was taken into consideration when estimating *in vivo* CL<sub>H</sub>.

### *In vitro* to *in vivo* predictions

*In vivo* CL_H_ in humans was predicted using lowest and highest predicted *in vivo* CL_int_ and measured *in vitro* fu for each drug and the well-stirred liver model (CL_H_ = (CL_int_ • f_u_ • Q_H_) / (CL_int_ • f_u_ + Q_H_), where Q_H_ is the liver blood flow). Predicted and observed *in vivo* CL_H_ were compared.

### *In silico* to *in vivo* predictions

The *in silico* methodology used in papers 3, 4 and 13 was used for *in silico* predictions of CL_int_, f_u_ and CL_H_. None of the drugs in the study was included in the training sets for the *in silico* models. Thus, every prediction was a forward-looking prediction where each compound was unknown to the models. Predicted and observed *in vivo* CL_H_ were compared.

## RESULTS & DISCUSSION

### *In vitro* based predictions

CL_H_ predicted from *in vitro* data differed up to 2436-fold (for gemfibrozil) depending on source of data (Table 2, Figure 1). For diclofenac, diazepam, naproxen and midazolam, ratios between highest and lowest predicted CL_H_ were 976-, 144-, 1634- and 39-fold, respectively (Table 2, Figure 1). Prediction errors ranged from 1.1-to 578-fold, with an average of 150-fold for lowest predicted estimates and 16-fold for highest predicted estimates. 70 % of the CL_H_-predictions had an error of ≤28-fold (n=20).

**Figure 1.**
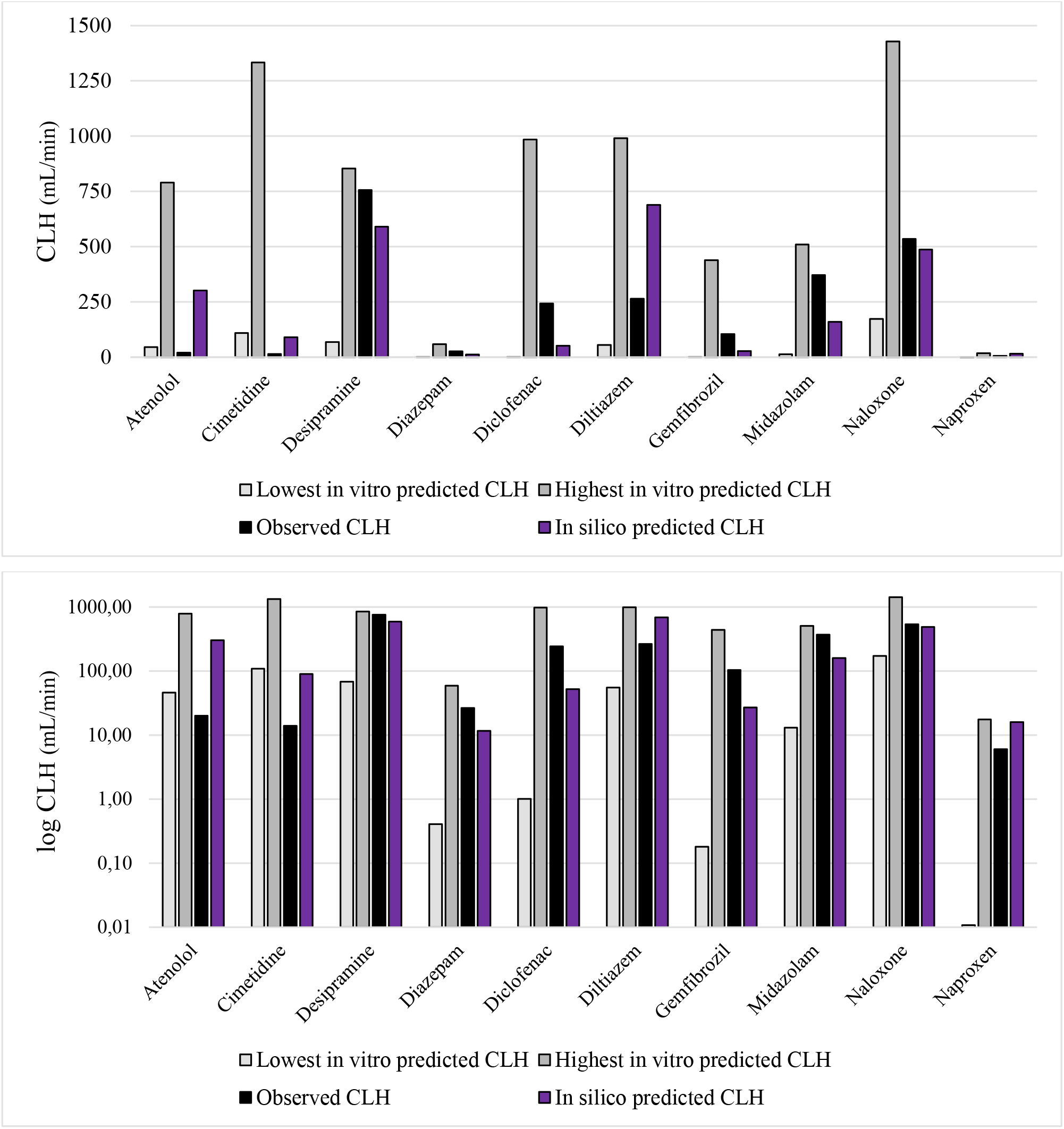
*In vitro* and *in silico* predicted and observed *in vivo* CL_H_ for the 10 drugs. a) linear CL_H_-scale, b) logarithmic CL_H_-scale.

For cimetidine, diclofenac, diltiazem and naloxone, predicted hepatic extraction ratios ranged from low to high depending of sources of CL_int_ and f_u_, and for cimetidine and naloxone, maximum predicted CL_H_ is incompatible with adequate/desirable PK-profile (too low predicted oral bioavailability). Minimum predicted CL_H_ of <1 mL/min for diazepam, gemfibrozil and naproxen (and potentially very long half-life) are also in line with risk of withdrawal during the preclinical phase.

Observed *in vivo* CL_H_ for all compounds except for atenolol and cimetidine were within the ranges for *in vitro* predicted CL_H_ (Figure 1).

Variability and reproducibility of these sizes could not only jeopardize compound drug selection and development, but also human safety in early clinical trials.

For atenolol and diclofenac, maximum prediction errors for CL_H_ obtained using *in vitro* data were 39- and 240-fold, respectively.

Atenolol (moderate permeability and gastrointestinal uptake, and efflux) has also shown highly variable Caco-2 permeability values (65-fold interlaboratory range and up to 16-fold intralaboratory variability) and efflux ratios (21-fold difference between laboratories and 4.4-fold intralaboratory variability, and 1/3 of measurements showing no efflux).^2,5,6^

Diclofenac (high permeability and complete gastrointestinal uptake) has reported S ranging between 2 (low) to 9000 (high) mg/L (7-30000 μM) and its half-life has been underpredicted by 5-fold and daily oral dose overpredicted by 70-fold based on laboratory data.^5,14^

### *In silico* based predictions

Mean, median and maximum prediction errors for *in silico* predictions of CL_H_ were 4.2-, 2.6- and 15-fold, respectively (Table 3, Figure 1). This is in line with results in other studies with our *in silico* prediction methodology.^3,4,8,12^ 70 % of CL_H_-predictions had maximum 4.7-fold prediction error. For both CL_int_ and f_u_ an approximate 5-fold 70 % confidence interval was predicted (70 % of observations/measurements anticipated to lie within ±5-fold errors; ANDROMEDA-based predictions).

**Table 3.**
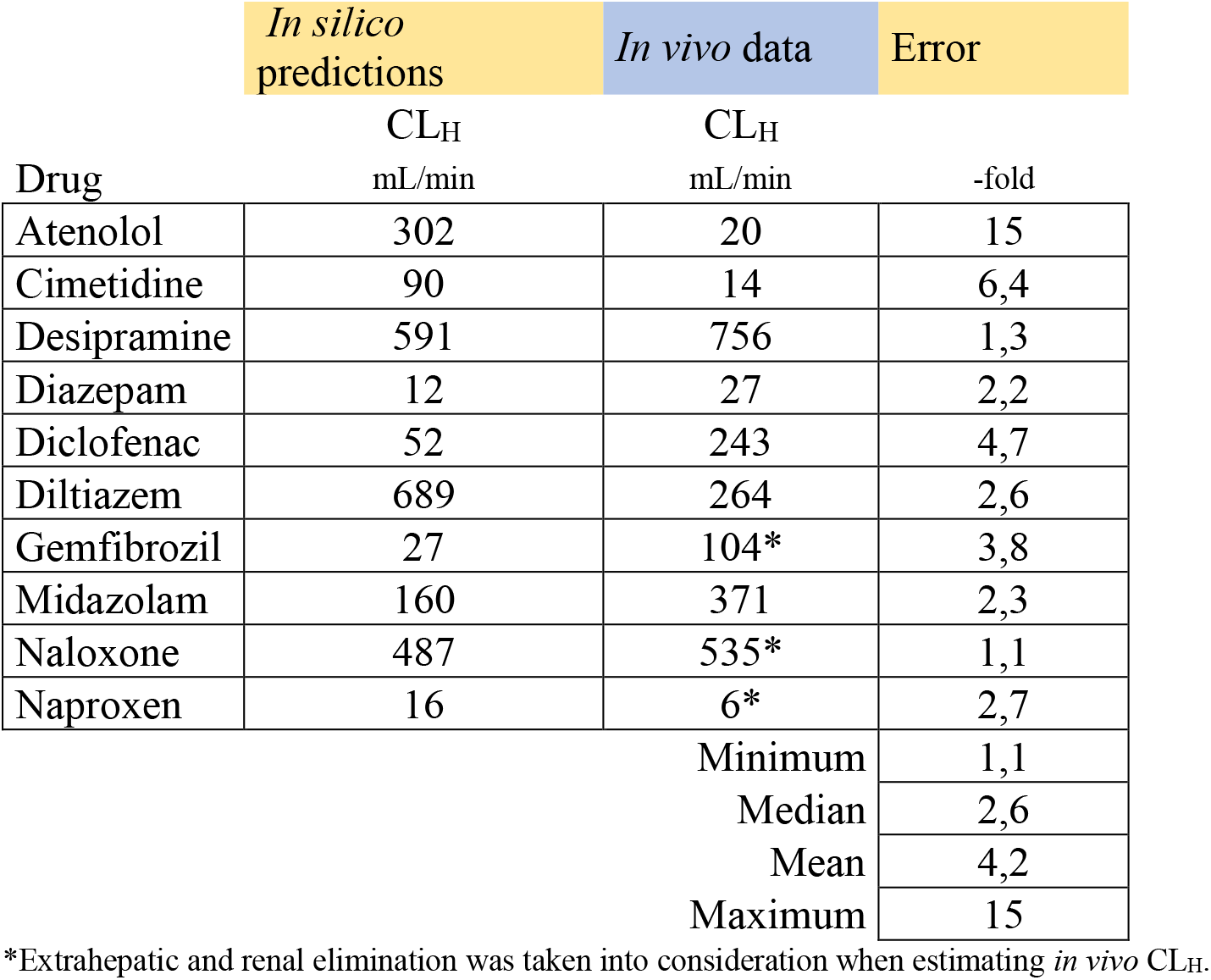
*In silico* predicted *in vivo* CL_H_ for the 10 selected drugs.

|  | *In silico*<br>predictions | *In vivo* data | Error |
| --- | --- | --- | --- |
| Drug | CL<sub>H</sub><br>mL/min | CL<sub>H</sub><br>mL/min | -fold |
| Atenolol | 302 | 20 | 15 |
| Cimetidine | 90 | 14 | 6,4 |
| Desipramine | 591 | 756 | 1,3 |
| Diazepam | 12 | 27 | 2,2 |
| Diclofenac | 52 | 243 | 4,7 |
| Diltiazem | 689 | 264 | 2,6 |
| Gemfibrozil | 27 | 104* | 3,8 |
| Midazolam | 160 | 371 | 2,3 |
| Naloxone | 487 | 535* | 1,1 |
| Naproxen | 16 | 6* | 2,7 |
|  |  | Minimum | 1,1 |
|  |  | Median | 2,6 |
|  |  | Mean | 4,2 |
|  |  | Maximum | 15 |

\*Extrahepatic and renal elimination was taken into consideration when estimating *in vivo* CL<sub>H</sub>.

There was no *in silico* prediction with high hepatic extraction ratio and corresponding low estimate (as found for *in vitro*-based predictions).

*In vitro*-based mean and maximum prediction errors for CL_H_ were 36- and 38-fold higher than corresponding *in silico* estimates, respectively.

Our *in silico* prediction methodology has previously been shown to be useful for accurate predictions of PK-parameters that have not been possible to quantify and predict using *in vitro* methods.^15^ With the new results it has also been successful for prediction of human PK for compounds with highly and uncertain *in vitro* PK.

## Conclusion

In conclusion, 36-to 38-fold higher average and maximum prediction errors of human CL_H_ were found for *in vitro* methodology compared to the *in silico* methodology. Mean and maximum *in silico* prediction errors were 4.2- and 15-fold, respectively, which is consistent with earlier results. The maximum prediction error found for the *in vitro* methodology was 578-fold. In contrast to the *in vitro* methodology the *in silico* models did not predict high hepatic extraction ratio for drugs with low CL_H_.

